# Three-dimensional genome reorganization during mouse spermatogenesis

**DOI:** 10.1101/585281

**Authors:** Zhengyu Luo, Xiaorong Wang, Ruoyu Wang, Jian Chen, Yusheng Chen, Qianlan Xu, Jun Cao, Xiaowen Gong, Ji Wu, Yungui Yang, Wenbo Li, Chunsheng Han, Fei Sun, Xiaoyuan Song

## Abstract

Three-dimensional genome organization plays an important role in many biological processes. Yet, how the genome is packaged at the molecular level during mammalian spermatogenesis remains unclear. Here, we performed Hi-C in seven sequential stages during mouse spermatogenesis. We found that topological associating domains (TADs) and chromatin loops underwent highly dynamic reorganization. They displayed clear existence in primitive type A spermatogonia, disappearance at pachytene stage, and reestablishment in spermatozoa. Surprisingly, even in the absence of TADs and chromatin loops at pachytene stage, CTCF remained bound at TAD boundary regions (identified in primitive type A spermatogonia). Additionally, many enhancers and promoters exhibited features of open chromatin and transcription remained active at pachytene stage. A/B compartmentalization and segmentation ratio were conserved in different stages of spermatogenesis in autosomes, although there were A/B compartment switching events correlated with gene activity changes. Intriguingly, A/B compartment structure on the X chromosome disappeared during pacSC, rST and eST stages. Together, our work uncovered a dynamic three-dimensional chromatin organization during mouse spermatogenesis and suggested that transcriptional regulation could be independent of TADs and chromatin loops at specific developmental stages.

## Introduction

Studies using open-ended Chromosome Conformation Capture (3C) methodology, Hi-C, have revealed fundamental insights into higher-order chromatin structure and three-dimensional (3D) genome organization in eukaryotes [1,2]. The higher-order chromatin is spatially packaged into a hierarchy of chromatin A/B compartments, topological associating domains (TADs), and chromatin loops [3,4]. Principal component analysis [1] (PCA) of Hi-C data uncovered that the genome is segmented into two types of compartments, where compartment A is associated with active chromatin regions while compartment B is associated with repressive chromatin regions. The switch between A/B compartments is related to transcriptional regulation and cell fate decision [5]. Although A/B compartments are pervasive and highly dynamic between different cells, the underlying mechanism of A/B compartment formation remains elusive. Unlike A/B compartments, which vary among cell types, TADs are largely invariant between different cell types and species except in the mitotic stage [3,6,7]. CTCF and cohesin, two key weavers of chromatin structure, were demonstrated to be highly enriched on TAD boundaries [3]. Recent studies reported that rapid degradation of CTCF or cohesin could eliminate TADs [8–10], suggesting that CTCF and cohesin are important for TADs formation and maintenance, while super-resolution chromatin tracing showed that TAD-like structures remained in a subset of single cells after cohesin depletion [11]. In addition to these 3C-based methods which are dependent on proximity ligation, a variety of emerging techniques based on different principles, such as GAM [12], SPRITE [13] and above mentioned super-resolution chromatin tracing [11], reinforced the existence of TAD-like structures in mammalian cells and even at single-cell level. Although chromatin architectures were widely assumed to tightly correlate with transcription control, recent findings cast several doubts on this assumption. On one hand, loss of TADs and loops impacts gene expression. For example, pathological disruption of TAD boundaries was reported to lead to improper gene activation in the adjacent TADs [14–16]. Also, enhancer-promoter loops were strongly affected by depletion of YY1 and correlate with gene deregulation [17]. Furthermore, Sven et al showed that transcription elongation could affect 3D genome organization [18]. On the other hand, some other studies suggest minimal role of transcription and 3D genome in modulating each other. For example, elimination of all TADs and chromatin loops by rapid degrading of cohesin complex had modest effect on gene transcription programs [9]. Many earlier reports found that transcription did not affect the establishment of chromatin organization during early development [19,20]. Thus, whether higher-order chromatin structures play a causative role in transcriptional regulation and whether it is dynamically regulated by transcription are unresolved and fundamental questions in molecular biology.

The mammalian spermatogenesis is a highly specialized developmental process which involves spermatogonia renewal and proliferation, meiosis, and spermiogenesis [21]. Spermatogenesis starts from a small number of primitive type A spermatogonia (PriSG-A), which can differentiate into type A spermatogonia (SG-A) and type B spermatogonia (SG-B) in sequence [22]. Subsequently, SG-B differentiate into preleptotene spermatocytes (plpSC), which undergo the final replication of nuclear DNA before entering meiotic prophase. In meiotic prophase, homologous recombination, including meiotic chromosome crossover (CO), occurs in pachytene spermatocytes (pasSC) [23]. Later on, the first cellular division takes place and produces secondary spermatocytes which rapidly and sequentially divide and form haploid round spermatids (rST), elongating spermatids (eST), and spermatozoa (SZ) [24]. Spermatogenesis is a complex and highly regulated process that is precisely controlled at both transcriptional, post-transcriptional, and translational levels [25,26]. Previous efforts elucidated the transcriptional regulation mechanisms of mammalian spermatogenesis from the aspects of transcriptome, DNA methylome and histone modifications [27–29]. Besides one dimensional (1D) information like histone modification and DNA methylation, important three-dimensional (3D) “epigenetic” information of the chromatin also needs to be properly “inherited” or “reprogrammed” during gamete generation. It thus constitutes a significant question to understand the organization and regulation of 3D chromatin structures during spermatogenesis.

In this study, we utilized versatile 3D genome and epigenome profiling methods to chart the landscape of the 3D genome organization and transcriptional regulation during mouse spermatogenesis. Our work provided new insights into the organization and dynamics of 3D chromatin structures in mammalian spermatogenesis.

## Results

### Chromatin TADs and loops were dynamically reorganized during mouse spermatogenesis

We systematically isolated eight spermatogenic cell types during mouse spermatogenesis by using the unit gravity sedimentation procedure [27,30]. These included primitive type A spermatogonia (PriSG-A), type A spermatogonia (SG-A), type B spermatogonia (SG-B), preleptotene spermatocytes (plpSC), pachytene spermatocytes (pacSC), round spermatids (rST), elongating spermatids (eST) and spermatozoa (SZ) (S1A Fig) [27,30]. The purities of these spermatogenic cell types (S1B Fig) were validated by morphological evaluation (S1A Fig), immunofluorescence staining (S1C Fig), and RT-qPCR (S1D Fig) with stage-specific markers.

Using different stages of spermatogenic cells, we conducted in situ Hi-C experiments [4], and generated high quality datasets as revealed by correlation analyses of biological replicates (S1 Table and S2A Fig). We obtained sufficient usable reads in all isolated cells except the SZ sample, so we included a published Hi-C data set of SZ cells in our analyses [20]. Square-shaped TADs are the most prominent architecture of the genome in Hi-C contact maps [3,31]. We first found that TADs underwent a drastic reorganization during spermatogenesis (Fig 1). The snapshots of our Hi-C contact maps (Fig 1) clearly showed that the triangle-shaped TADs were well detected in PriSG-A, SG-A and SG-B, but became much weaker in plpSC. TADs completely disappeared at pacSC and rST stages, and later started to re-emerge in eST and ultimately were reestablished in SZ. To confirm this reorganization pattern on a genome-wide basis, we performed a meta-domain analysis. We aggregated chromatin interaction maps of all TADs identified in PriSG-A, which clearly recapitulated the patterns observed by manual inspection, i. e. TADs were lost in pacSC and re-established in eST and SZ (Fig 2A). To quantitatively gauge the dynamics of TADs, we utilized the “insulation score (IS)” algorithm [32] to measure the strength of TAD boundaries in different stages. This score reflects the strength of TAD boundaries as calculated by the chromatin interaction frequency between the upstream and downstream regions of each genomic locus. We found that the insulation scores diminished gradually from PriSG-A to pacSC (Fig 2B). Unlike most previous findings that TADs appeared very stable and invariant across cell types or even species [3,6], our Hi-C maps strikingly revealed that TADs were distinctly and dynamically reorganized during spermatogenesis (Fig 1 and Fig 2). Apart from TADs level, we examined whether chromatin loops [4] (focal interaction points in the Hi-C map) underwent a similar reorganization during spermatogenesis. We used one of the very stringent loop calling algorithms, HICCUPS [4,33], to identify bona fide chromatin loops in the PriSG-A sample. We then used these identified loops to perform Aggregation Peak Analysis (APA) [4] in order to characterize the loop strength. In accordance with the meta-domain analysis, the meta-loop plot of APA revealed massive reorganization of chromatin loops (Fig 3A), which could be quantitatively reflected by the APA P2LL values (the intensity ratio between the central pixel and the mean of the pixels in the lower left corner, representing the strength of chromatin loops). P2LL values of chromatin loops strongly decreased from PriSG-A to pacSC, but returned to a certain level in SZ (Fig 3B). These results suggested that both TADs and chromatin loops reorganized dramatically during spermatogenesis, and their reorganization patterns strongly concurred with each other.

**Fig 1.**
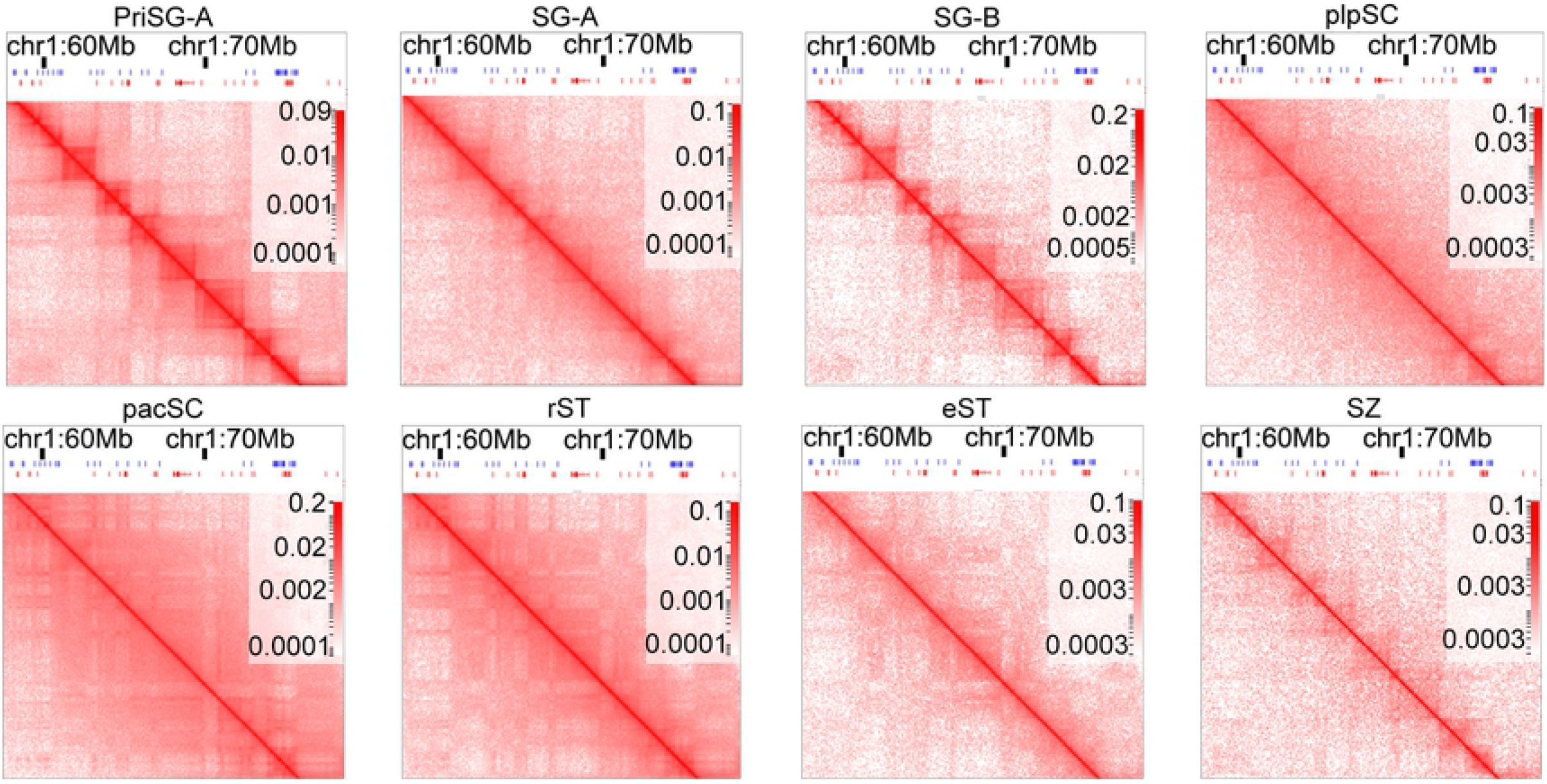
An overview of chromatin domains (TADs) reorganization during mouse spermatogenesis. Snapshots of Hi-C ICE (iteratively corrected) normalized contact maps at a region on chromosome 1 (chr1: 60Mb to 70Mb) at eight different stages (PriSG-A, SG-A, SG-B, plpSC, pacSC, rST, eST and SZ) during mouse spermatogenesis. These snapshots clearly showed that triangle shaped structures-TADs existed in PriSG-A, SG-A, and SG-B cells, started to disappear from plpSC stage to eST stage, and re-emerged in SZ. It also should be noticed that the plaid pattern of Hi-C maps indicated that compartment structures existed at all stages.

**Fig 2.**
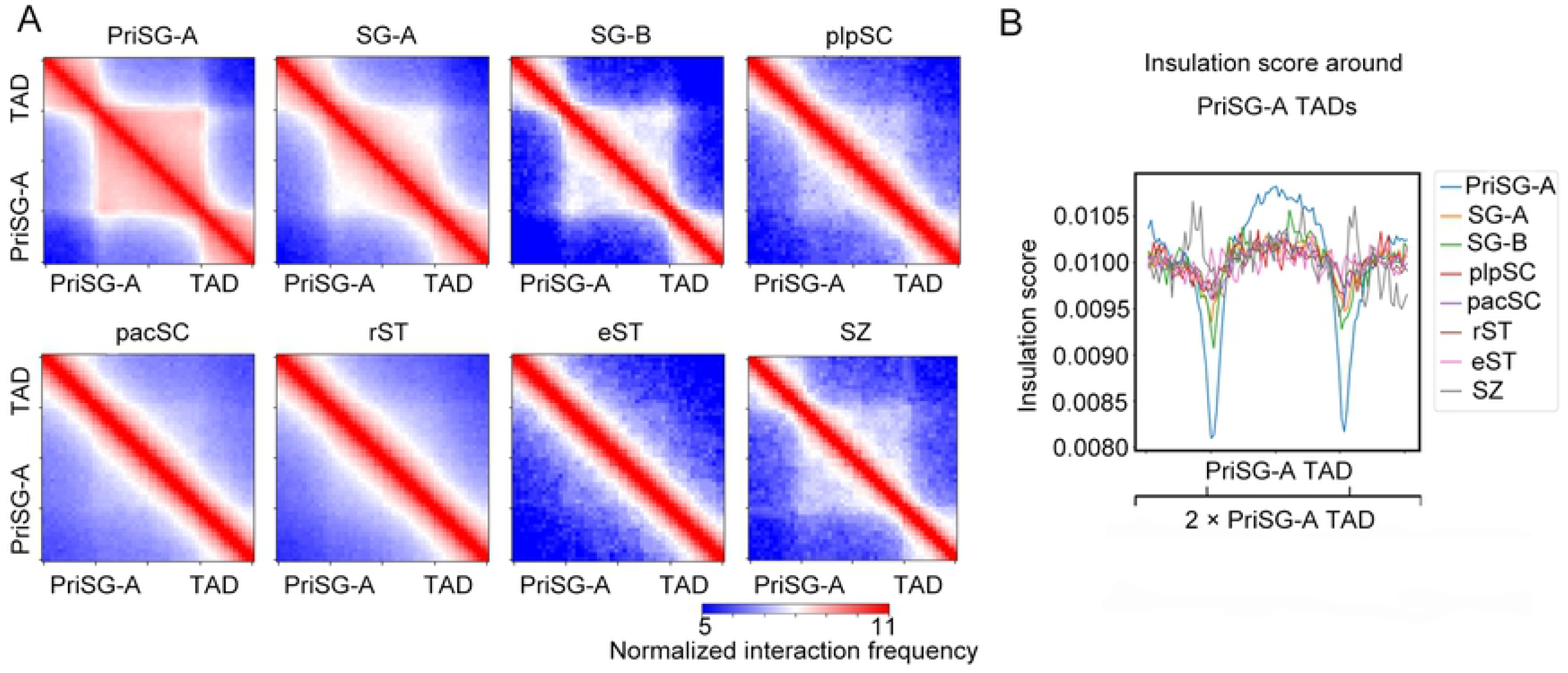
Reorganization of TADs during mouse spermatogenesis. (A) ICE normalized chromatin interaction maps around every TAD identified in PriSG-A were piled up and formed aggregated Hi-C heatmaps, showing the normalized average interaction frequencies for all TADs (defined in PriSG-A) and their nearby regions (±0.5 TAD length) in PriSG-A, SG-A, SG-B, plpSC, pacSC, rST, eST and SZ stages during mouse spermatogenesis. Heatmap scales represent the averaged and ICE normalized Hi-C reads numbers. (B) The average insulation scores plotted around TAD boundaries (defined in PriSG-A) in the above eight different stages during mouse spermatogenesis.

**Fig 3.**
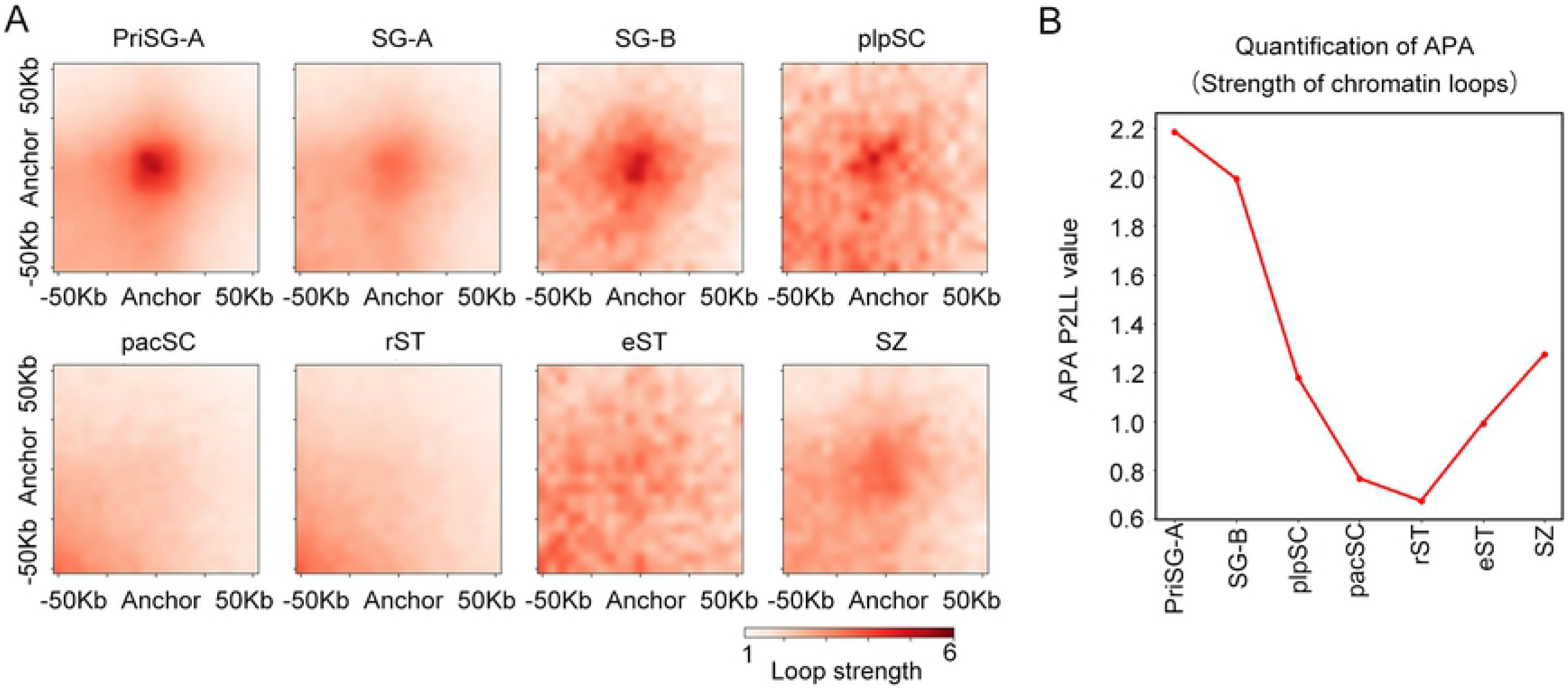
Chromatin Loops were reorganized during mouse spermatogenesis. (A) The APA (Aggregation Peak Analysis) heatmaps of all KR normalized [34] Hi-C interactions near high-confidence chromatin loop regions (Upstream / Downstream 50 kb of loop anchors) identified by HICCUPS (see Methods) in eight stages (PriSG-A, SG-A, SG-B, plpSC, pacSC, rST, eST and SZ). The peak at the center of the APA plot indicated the aggregated signal from our peaks set as a whole. (B) The APA Peak to Lower Left (P2LL) values (the ratio of the central pixel to the mean of the pixels in the lower left corner), quantitatively characterizing the dynamics of chromatin loops strength across different spermatogenesis stages.

### Chromatin remained accessible in transcriptionally active pacSC in the absence of TADs and chromatin loops

Genome-wide analyses of our PriSG-A Hi-C data identified 2,045 TADs and 2,435 high-confidence chromatin loops, which were comparable to previous Hi-C studies in mouse cell lines [3,4]. By contrast, only less than 100 TADs and loops could be detected in pacSC (Fig 4A and 4B), reinforcing the conclusion that most of the chromatin TADs and loops were lost in pacSC. With such a remarkable reduction of TADs and chromatin loops, we wondered how the 3D genome correlated with 1D regulatory landscape in pacSC. High-quality reproducible ATAC-seq datasets were acquired on four different types of cells (PriSG-A, SG-A, pacSC, and rST) to deduce the global chromatin accessibility (S2B Fig). Most accessible chromatin regions were located at gene promoters and enhancers, and the chromatin accessibility remained largely invariant as spermatogenesis proceeded (Fig 4C, S2C Fig and S2D Fig). By comparing PriSG-A and pacSC, we identified 6580 (out of 15815) differentially accessible chromatin regions; while between pacSC and rST, there were 1288 (out of 15815) differentially accessible regions (S3A Fig). This relatively mild change of chromatin opening contrasted strongly with the dramatic loss of chromatin 3D organization in pacSC (TADs and loops, Figs 1, 2 and 3). We showed one of the representative regions on chromosome 12 to demonstrate this striking contrast in pacSC as compared to PriSG-A (Fig. 4C). These results suggested an independent relationship between chromatin looping and chromatin accessibility. Furthermore, ChIP-Seq of serine-2-phosphorylated RNA Polymerase II (S2P, an elongating form of RNA Pol II) showed that open chromatin regions were actively transcribed at pacSC and rST stages (Fig 4D). Despite being infrequent, the relatively small percentage of chromatin regions displaying altered ATAC-seq peak intensities between PriSG-A and pacSC were highly associated with meiotic recombination and/or recombination-relevant processes, as revealed by functional enrichment analysis (S3B Fig). The regions with differential chromatin accessibility between pacSC and rST were mainly associated with meiosis (S3C Fig). This result was consistent with the fact that meiotic homologous recombination occurs in pacSC, and rST was produced after meiosis was accomplished. Our analyses of the transcription factor (TF) motifs enriched in the subset of differential chromatin accessibility revealed both reported key TFs and unknown TFs, orchestrating the transitions of cell identities during spermatogenesis (S3D, S3E, S3F and S3G Fig). For example, the chromatin regions closed in PriSG-A but opened in pacSC were enriched with TF motifs like A-MYB and RFX2 (S3D Fig), which were well-known master TFs for male meiosis [35,36], and with some newly identified TFs such as ELK4 (S3D Fig). The chromatin regions opened in PriSG-A but closed in pacSC were enriched with TF motifs for known PriSG-A regulators such as SP1 and DMRT1/6 (S3E Fig) [37,38], and also for unknown TFs like KLF5 and KLF14 (S3E Fig). The sites of meiotic DNA double strand breaks (DSBs) have been reported to correlate with chromatin accessibility [39]. We thus compared the numbers of accessible meiotic DSB sites (identified by a published Dmc1 ChIP-Seq [40]) between PriSG-A and pacSC. Unexpectedly, we found that the numbers of the accessible DSB sites were largely unchanged (S3H Fig), suggesting that meiotic DSB sites were pre-opened prior to the pacSC developmental stage. As pacSC piRNAs are unique transcription units that are highly expressed at pacSC stage and are required for the followed stages of spermatogenesis [41], we also compared the numbers of accessible piRNA clusters between PriSG-A and pacSC. A larger number of accessible piRNA clusters were found in pacSC but not in PriSG-A (S3I Fig), which was consistent with the fact that pacSC transcribes pacSC-specific piRNAs. These data together revealed a relatively stable chromatin accessibility landscape during spermatogenesis, although most TADs and chromatin loops did not exist in pacSC. There were relatively mild changes in specific regions that correlated with known transcriptional activity during spermatogenesis.

**Fig 4.**
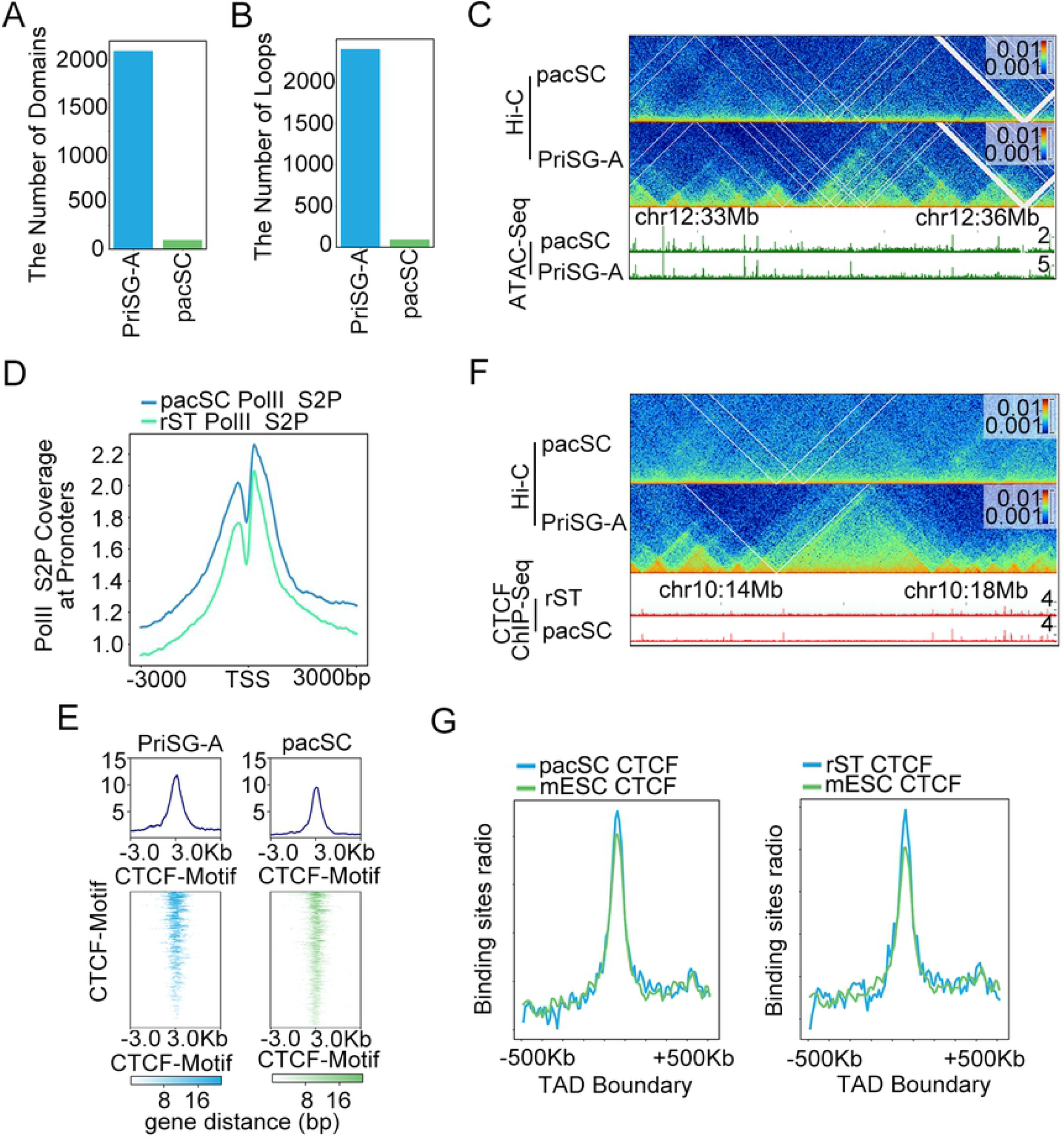
Loops and TADs were disappeared at pacSC stage while the chromatin accessibility of CTCF binding sites and CTCF binding on chromatin were not significantly changed. (A) The numbers of the identified TADs in PriSG-A and pacSC. (B) The numbers of the identified chromatin loops in PriSG-A and pacSC. (C) Chromatin structures and accessibility of a region on chromosome 12. Hi-C heatmaps showed the dramatic difference of TADs (top panel) at pacSC and PriSG-A stages, and ATAC-seq coverage tracks showed the unchanged chromatin accessibilities of the same regions (bottom panel) at pacSC and PriSG-A stages. (D) The meta-gene coverage of RNA Pol II S2P ChIP-Seq in pacSC and rST showed that RNA Pol II S2P were still actively bound at TSS regions. (E) The ATAC-seq meta-gene plot around CTCF motif regions suggested that chromatin was still open at CTCF binding sites during pacSC stage. (F) Hi-C contact maps and CTCF ChIP-Seq tracks of a region on chromosome 10. Hi-C heatmaps showed that TADs disappeared in pacSC comparing to PriSG-A (top panel), and CTCF ChIP-Seq peaks (bottom panel) at pacSC and rST stages showed that CTCF still bound at TAD boundaries (defined in PriSG-A). (G) The binding sites distribution plots of CTCF in pacSC (left), rST (right) and mESC (ENCODE mouse CTCF ChIP-Seq data) showed that CTCF binding was highly enriched at TAD boundary regions (defined in PriSG-A) in pacSC and rST stages.

### CTCF binding remained unchanged when TADs disappeared at pacSC stage

With the ATAC-seq data, we could infer the TF occupancy at the TF motif regions from the chromatin accessibility [42]. We particularly focused on the CTCF motif as it was one of the major weavers of 3D chromatin structure [43]. Interestingly, we found that chromatin accessibilities at CTCF motifs appeared highly similar between PriSG-A and pacSC (Fig 4E). This result suggested that CTCF could still bind to chromatin in cells without TADs (i.e. in pacSC and rST). To further confirm this possibility, we performed CTCF ChIP-Seq in pacSC and rST. Our data showed that CTCF bound to TAD boundaries (according to those in PriSG-A) although TADs were largely lost in pacSC and rST (Fig 4F and 4G). The binding of CTCF exhibited a comparable strength as that in mESC cells from ENCODE data (Fig 4G). These findings suggested that CTCF binding to chromatin was not sufficient to maintain chromatin loops and TADs, and that the CTCF binding on the 1D genome (by ChIP-Seq or motifs) did not predict the 3D connections mediated by CTCF (CTCF looping).

### The dynamic reorganization of A/B Compartments was independent of TADs during spermatogenesis and showed difference in autosomes and sex chromosome

The 3D genome consists of loops, TADs and A/B compartments [44]. The dramatic changes of TADs and chromatin loops during spermatogenesis led us to examine the dynamics of the compartments. We found that the A/B compartmentalization pattern (~40% A versus ~60% B) on autosomes were largely unaltered at eight different stages of spermatogenesis (Fig 5A and S4A Fig), suggesting that the chromatin structure represented by compartments was independent of TADs and loops. Compartment strength for separating A and B compartments, as measured by absolute PC1 value in different stages, showed that SG-B had the highest compartment strength while pacSC had the lowest (Fig 5B). We further conducted saddle strength analysis [45] to better quantify the compartment strength at the eight stages. By comparing saddle plots and saddle strength scores (AA + BB)/2*AB (Fig 5C and S4B Fig), we found that SG-B had the highest and pacSC had the lowest compartment strength. This was consistent with saddle analysis, suggesting that the higher-order chromatin was less compartmentalized in pacSC. By calculating the correlation of PC1 values (representing compartment strength), we found that a majority of the compartments was consistent between different stages (S5A Fig). However, the PCA analysis based on compartment scores (PC1 values) (S5B Fig) revealed two distinct clusters of the eight stages (Fig 5D), suggesting that the switch between A/B compartments happened frequently. Quantitatively, we found that around 30 percent of the regions underwent compartment switching during spermatogenesis. After counting the sequential compartmental switches between two stages, we found that the numbers of A to B and B to A switches varied during the whole process of spermatogenesis and the numbers of switching compartments rarely exceeded 10 percent of all compartments in the genome (S5C Fig). Given the fact that A/B compartments largely correlated with the active/repressive chromatin status, the dynamic switching of A/B compartments would permit the dynamic transcriptional regulation of spermatogenesis. As an example, we examined the *Kit* gene locus. The *Kit* gene is specifically expressed in SG-A and is required for the differentiation of spermatogonia [46,47]. It was located in the active A compartment in PriSG-A and SG-A but converted to an inactive B compartment in pacSC (S5D Fig), accompanying the shut-down of the *Kit* gene expression.

**Fig 5.**
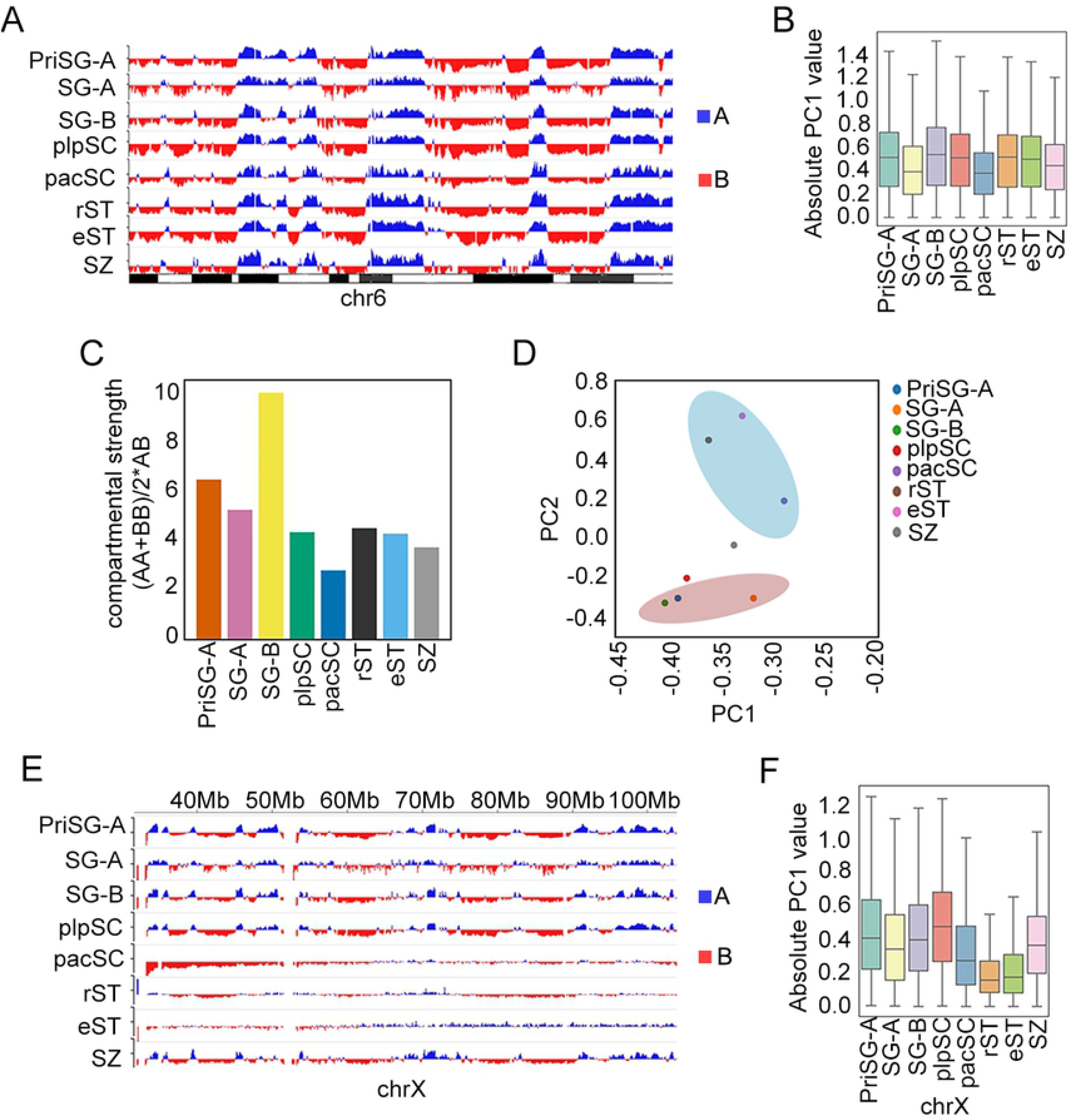
Dynamics of A/B compartments during spermatogenesis. (A) Compartment eigenvector was plotted along the linear sequence of a region on chromosome 6 for eight different spermatogenetic stages and used as A (blue) and B (red) compartments segmentation. (B) The absolute PC1 values (representing compartment strength) of eight different stages during mouse spermatogenesis on autosomes. (C) Compartment strength plots: ratios of corner interaction scores (AA + BB) /(AB +BA), where the saddle data were grouped over the AA+BB corners and AB+BA corners, characterizing the intra-compartment interactions versus inter-compartment interactions [45]. (D) The PCA analysis based on compartment scores of eight different stages during mouse spermatogenesis. (E) Compartment eigenvector was plotted along the linear sequence of chromosome X for eight different spermatogenetic stages and used as A (blue) and B (red) compartments segmentation. (F) The absolute PC1 values (representing compartment strength) of eight different stages during mouse spermatogenesis on chromosome X.

In addition, we interrogated the compartmentalization of the X chromosome. PCA analysis of X chromosome Hi-C data at pacSC, rST, and eST stages did not show clear segmentation of the X chromosomes (Fig 5E). By contrast, these three stages had low absolute PC1 values of X chromosomes, indicating low compartment strength (Fig 5F). The loss of A/B compartments on X chromosome during these three stages might be due to the fact that X chromosome underwent meiotic sex chromosome inactivation (MSCI) during meiosis [48]. Since the Hi-C reads coverage on the Y chromosome was limited to the first 3Mb of the total 16 Mb size (data not shown), we did not do further analysis on the Y chromosome.

### Meiotic pacSC chromatin structure was distinct from mitotic chromatin structure

To gain an overview of the biophysical feature of the 3D genome organization during spermatogenesis, we performed *P(s)* analyses, which depicted the contact frequency of any genomic region in Hi-C data as a function of the linear distance between different regions. Hi-C maps are characterized by a general decay of contact frequency P with genomic distances [1]. The *P(s)* curves of different spermatogenic stages revealed versatile chromatin folding patterns with two extreme cases - pacSC and SZ (Fig 6A). In the first-three stages-PriSG-A, SG-A and SG-B, the *P(s)* curves were highly consistent with previous reports that the interphase cells generally followed the rule of *P(s)* ~ s (slope) as −1 [1,49], suggesting that the chromatin structures of PriSG-A, SG-A and SG-B were similar to that in other interphase cells (Fig 6B). Starting from plpSC stage, the P(s) curve’s slope began to increase, and reached a rather flattened slope (~ −0.5) (Fig 6C). The *P(s)* curves of pacSC and plpSC showed that the slopes varied differently at different genomic distances, and there was a steep drop at ~ 8Mb distance in the *P(s)* curve of pacSC, revealing a special chromatin structure and folding pattern comparing to that in PriSG-A (Fig 6C). It has been reported that mitotic chromatin displayed different folding pattern as compared to that of interphase chromatin [45,49], As both meiotic cells in pachytene and mitotic cells undergo chromosome condensation, we compared the 3D chromatin structures in mouse meiosis (pacSC) versus mitosis. The *P(s)* curves showed that within 1Mb genomic distance, mitotic and meiotic chromatin shared similar folding patterns with *P(s)* ~ s (slope) as −0.5, but they had a striking difference at a longer distance, where the meiotic curve had a steep drop at ~ 8Mb while the mitotic curve dropped at ~ 30Mb distance (Fig 6D). In addition, our current work reveled that meiotic cells clearly contain A/B compartments (Fig 5A and S4A Fig), contrasting with mitotic chromatin which was reported to lack such structure [49]. Together our data revealed many distinctive 3D features of the meiotic pacSC chromatin versus the mitotic chromatin. Subsequent to the pacSC stage in development, the *P(s)* curve continued to change. At the distance smaller than 30Mb, the *P(s)* of two certain chromatin regions tended to gradually decrease from pacSC to SZ, while *P(s)* increased at the distance greater than 30Mb (Fig 6A). In general, our *P(s)* analysis clearly described the dynamic 3D genome reorganization during spermatogenesis.

**Fig 6.**
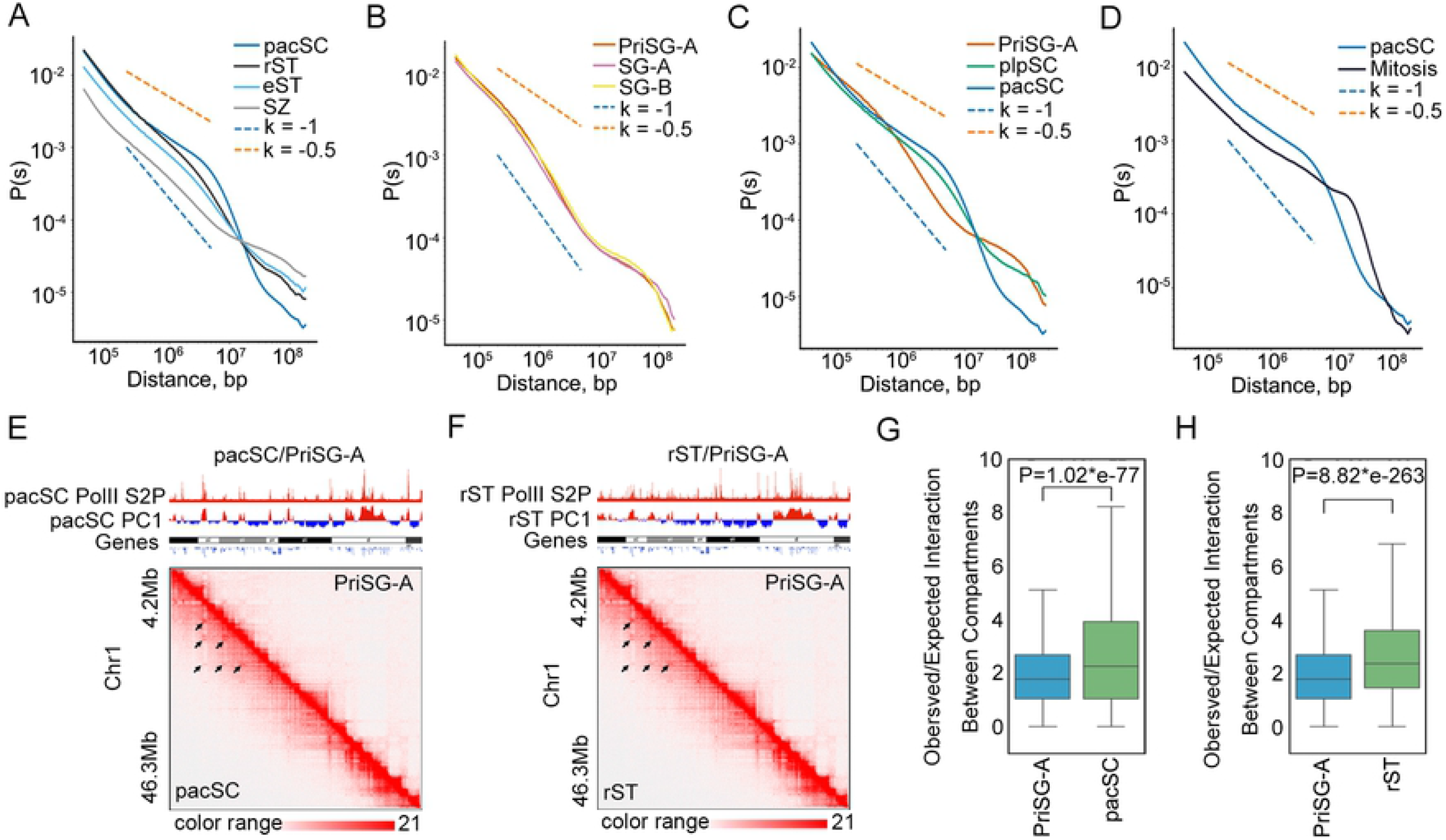
Distinct chromatin structures during mouse spermatogenesis and the transcriptionally correlated compartments. (A) The P(s) curves (relationship between interaction probability and genomic distance) of pacSC, rST, eST and SZ, and slopes (k) −1 (orange) and −0.5 (blue) were shown by the dotted lines. (B) The P(s) curves (relationship between interaction probability and genomic distance) of PriSG-A, SG-A and SG-B, and slopes (k) −1 (orange) and −0.5 (blue) were shown by the dotted lines. (C) The P(s) curves (relationship between interaction probability and genomic distance) of PriSG-A, plpSC and pacSC, and slopes (k) −1 (orange) and −0.5 (blue) were shown by the dotted lines. (D) The P(s) curves (relationship between interaction probability and genomic distance) of pacSC and mitotic cells, and slopes (k) −1 (orange) and −0.5 (blue) were shown by the dotted lined. (E and F) Snapshots of transcriptionally correlated compartments at pacSC stage (E) or rST stage (F) compared to that in PriSG-A. Upper panels showed the Pol II S2P ChIP-Seq peaks, PC1 values which stand for A/B compartments and genes (from top to bottom). Lower panels were the Hi-C maps (the upper right corner was PriSG-A stage while the bottom left corner was pacSC stage in E or rST stage in F). The black arrows pointed to the transcriptionally correlated compartments. (G and H) The boxplot of local (no longer than 10Mb) top A to top A compartment interactions (observed/expected), showing that A to A compartment interactions enhanced specifically at pacSC (G) and rST (H) stages.

### The local A to A compartment interactions were enhanced at pacSC/rST stages and were highly correlated with RNA Pol II transcription

Our manual inspection of the Hi-C maps revealed locally enhanced large-scale A to A compartment interactions in pacSC and rST compared to that in PriSG-A, although genome-wide compartmentalization in pacSC and rST were decreased (Fig 6E and 6F). This result was consistent with a previous study showing that the interactions between certain compartments were strengthened when TADs diminished upon rapid degradation of cohesin [9]. To elaborate this phenomenon, we calculated the local Hi-C interaction frequencies (distance no longer than 10Mb) between the strongest A compartments (the top 300 compartments ranked by PC1 values in PCA analysis), and found that these strongest A-A compartment interactions were largely strengthened in both pacSC and rST, comparing to that in PriSG-A (P <0.05, Man-Whitney U test) (Fig 6G and 6H). These pacSC and rST specific enhanced A to A compartment structures were highly enriched for transcriptional activities as indicated by our Pol II S2P ChIP-Seq (Fig 6E and 6F), suggesting that these enhanced compartments structures were transcriptionally correlated and potentially driven by local transcriptional hyper-activities at these stages.

## Discussion

The 3D genome organization during meiosis is poorly characterized. It remains dogmatic that the hierarchical structures of the genome, especially TADs, are rather static among cell types or even among species. Mouse spermatogenesis is a perfect model system for investigating the dynamics of chromatin organization, as there are sequential, step-wise progressions of chromatin landscapes, and highly regulated gene transcription programs in this process, which are also accompanied by dramatic chromatin condensation in both mitosis and meiosis.

Here, we conducted systematic epigenomic profiling and Hi-C analysis in eight sequential spermatogenic cells/stages and documented highly dynamic reorganization of the 3D genome architecture during mouse spermatogenesis. These datasets revealed that the higher-order chromatin structures underwent dramatic changes during spermatogenesis in the scales of TADs and chromatin loops (Fig 7). The TADs and chromatin loops disappeared in pacSC during meiosis and reestablished in SZ. Our results showed that in pacSC, when TADs were absent, promoters and enhancers remained open, the compartments were preserved, and RNA Pol II binding increased at A compartment regions. These results indicated that transcription could be independent of higher-order chromatin organization on the scales of TADs and chromatin loops (but was related to compartments) at specific developmental stages. Our finding of the independence between compartments and TADs or chromatin loops was consistent with recent reports that after loss or knockdown of CTCF, the TADs and chromatin loops were disrupted while the compartments were still stable [8]. However, we also found that CTCF was still bound to TAD boundary regions even when TADs disappeared, demonstrating that CTCF itself was not sufficient to maintain TADs. Reminiscently, one previous study showed that rapid degradation of CTCF had limited effects on TADs, suggesting CTCF itself is also not necessary to maintain TADs [50]. The bona fide role of CTCF in 3D genome organizations warrant further characterizations. Also, future efforts on identifying additional factors regulating TADs will broaden our understanding on the function and mechanism of higher-order genome organization.

**Fig 7.**
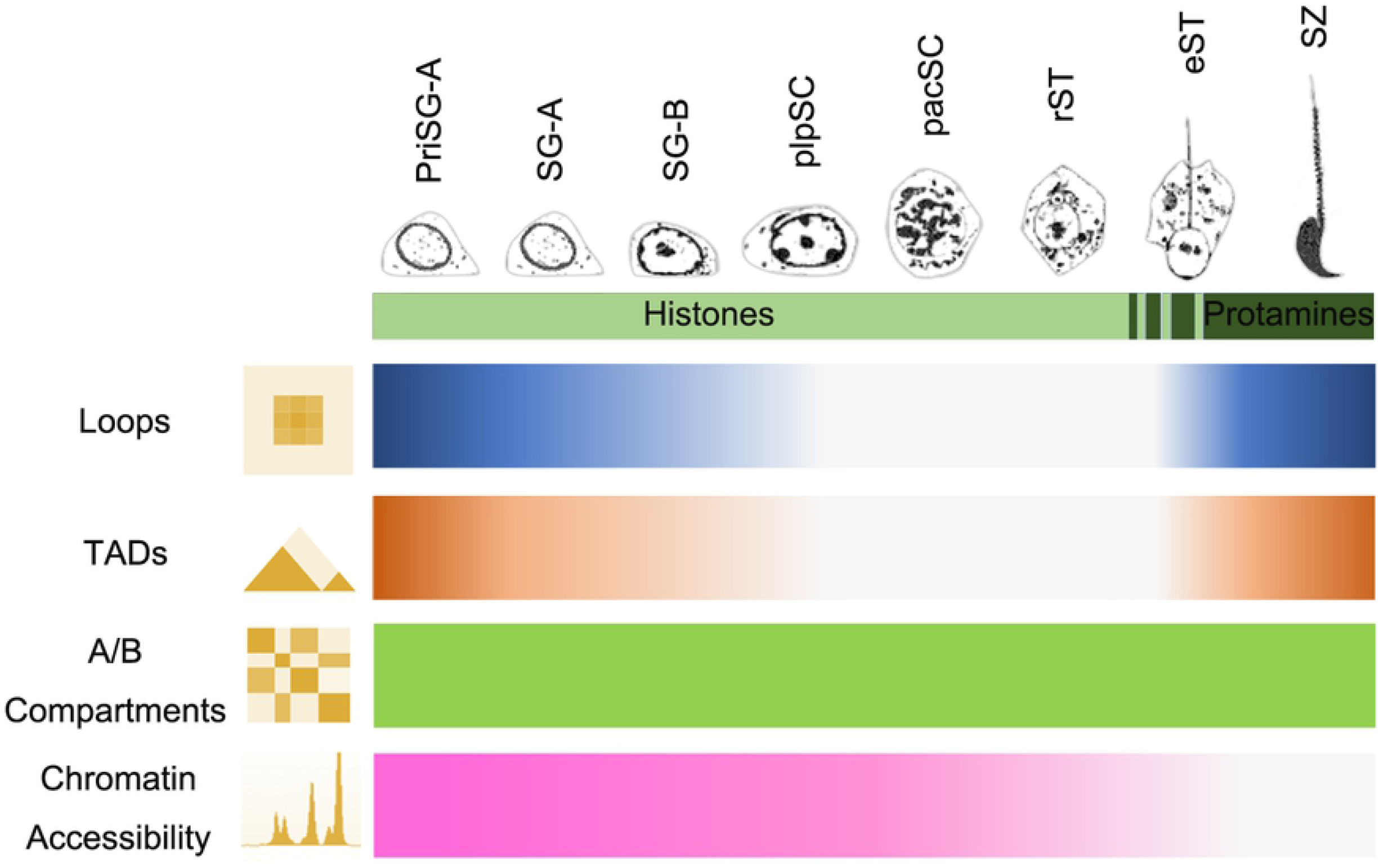
Diagram of 3D genome organization and chromatin accessibility during mouse spermatogenesis. Chromatin loops and TADs were reorganized dynamically during mouse spermatogenesis, where they clearly existed in PrisSG-A, SG-A and SG-B, started to disappear in plpSC, and totally disappeared in pacSC and rST, and then started to re-establish in eST and finally reestablished in the sperm. A/B compartments existed during the whole process from PriSG-A to sperm on autosomes. The chromatin was accessible in promoters and enhancers during the whole spermatogenesis process, although the accessibilities were gradually decreased.

Finally, the strengthened transcriptionally correlated compartments in pacSC with extremely weakened TADs and chromatin loops suggested that transcription might be independent with TADs and chromatin loops but still related to compartments. How compartments contribute to gene expression in this process awaits future investigations. Taken together, our study not only provided a comprehensive view of chromatin organization dynamics during mammalian gametogenesis, but also yielded novel insights on the fundamental relationship between 3D genome and transcription.

This is the first-ever study of 3D chromatin reprogramming during meiosis in mammals, together and independent with the other three groups that just published the similar work within one month in *Molecular Cell* (Wang et al., 2019, DOI:10.1016/j.molcel.2018.11.019) and *Nature Structural & Molecular Biology* (Alavattam et al., 2019, DOI: 10.1038/s41594-019-0189-y; Patel et al., 2019, DOI: 10.1038/s41594-019-0187-0). Our major findings were well validated by results from the other three papers. These include: 1) TADs were depleted in pacSC (showed in all papers). 2) Conventional A/B compartments were attenuated in pacSC accompanied by emergence of transcriptionally related compartments (between A-A compartments) in pacSC. 3) Meiotic genome folding shared similarity with that of mitotic genome, yet they were different. There were also some things that our paper had while the others did not: 1) We have characterized more comprehensive and fine profiling of the 3D genome during spermatogenesis (8 stages in total); 2) Our manuscript included not only 3D organization of the genome and gene expression, but also datasets for chromatin accessibility. 3) We performed RNA Pol II S2P and CTCF-ChIP-Seq in pacSC stage, and the data suggested that CTCF was not sufficient to create TAD boundaries and the independence between transcription and TADs.

## Materials and Methods

### Experimental Animals

C57BL/6 mice were housed in the Animal Center, University of Science and Technology of China, and cultured under a 12-h light/dark cycle (lights off at 7 p.m.) at 23±2°C. All animal manipulations were conducted in strict accordance with the guidelines and regulations set forth by the University of Science and Technology of China (USTC) Animal Resources Center and University Animal Care and Use Committee. The protocol was approved by the Committee on the Ethics of Animal Experiments of the USTC (Permit Number: PXHG-SXY201510183) for mouse experiments.

### Purification of male germ cells during spermatogenesis

C57BL/6 mice were originally purchased from Vital River Laboratories in Beijing, China. Primitive type A spermatogonia (PriSG-A), type A spermatogonia (SG-A), type B spermatogonia (SG-B), preleptotene spermatocytes (plpSC), pachytene spermatocytes (pacSC), round spermatids (rST) and elongating spermatids (eST) were isolated by the unit gravity sedimentation procedure based on Bellvé and Gan’s description[27,30] with minor modification. PriSG-A was isolated from 6-days postpartum (dpp) mice; SG-A and SG-B were isolated from 8-dpp mice; plpSC was isolated from 17-dpp mice; pacSC, rST and eST were isolated from adult mice. In brief, the testes were removed and minced by scissors until a semiliquid state had been achieved. Then this tissue was incubated in 10 ml DMEM (Gibco, 11995-081) containing 1 mg/ml collagenase IV (Sigma, C5138-1G) and 1 unit/ml DNase I (Sigma, AMPD1-1KT) in a shaking water bath at 37 °C for 5 min. 20 ml of fresh DMEM containing 10% FBS was added to stop the digestion. The seminiferous tubules were collected by centrifugation at 500 g for 2 min. The pellet was washed once with DMEM and was resuspended in 10 ml DMEM containing 1 mg/ml Trypsin (Sigma, T1426-500MG) and 1 unit/ml DNase I (Sigma, AMPD1-1KT), and incubated in a shaking water bath at 37 °C for 5 min. 20 ml of fresh DMEM containing 10% FBS was added to stop the digestion. Cells were collected by centrifugation at 500 g for 2 min. The cell pellet was washed twice with DMEM and resuspended in 40 ml DMEM containing 0.5% BSA (Sangon, AD0023-100g). Then the cells were filtered through a 40 μm Nylon Cell Strainer (BD Falcon, 352340) and separated by sedimentation velocity at unit gravity at 4 °C, using a 2-4% BSA gradient in DMEM. Only fractions with expected cell type and purity (≥ 75%) were pooled together. The collected cells were then cultured in 10 ml DMEM containing 10% FBS in a 10 cm diameter tissue culture dish precoated with 0.1 mg/ml poly-D-lysine for 3 hours at 34 °C. Sertoli cells (SE) attached to the culture plates, and the germ cells in suspension were collected by centrifugation at 500 g for 5 min. Spermatozoa (SZ) were isolated from the adult mice, too. The cauda epididymis were removed and cross cut in a tissue culture dish containing PBS (Thermo, 21600-010) preheated at 37 °C. After a while, SZ swam up. The motile sperm were collected and washed twice with PBS. Then SZ were filtered through a 40 μm Nylon Cell Strainer and collected by centrifugation at 500 g for 5 min. The purity of SZ was evaluated at ~95% based on morphological characterization. And the purities of the other isolated cells were evaluated and identified by their morphological characterization, immunofluorescence staining and RT-qPCR with germ cell type-specific markers (for example, GFRa1 for PriSG-A, Kit for SG-A, SCP3 and γH2A.X for pacSC, CLGN for rST and WT1 for SE.).

### Immunofluorescence

Immunofluorescence analysis was performed to identify the cell type and purity of the isolated cells. Since the isolation methods were routine for us with high purity and success [27], only part of the isolated cell types were used to perform Immunofluorescence. Specially, the cell pellet was resuspended in 5 μl FBS and transferred to glass slides precoated with 1% gelatin. 10 min later, cells were fixed with 4% paraformaldehyde (Sigma, V900894-100G) for 30 min and washed three times with PBS, permeabilized with 0.5% Triton X-100 (Sigma, 93443-100ML)/PBS for 10 min and washed three times with PBS, and blocked with 3% BSA/PBS for 1 hour. Then the cells were incubated with the primary antibodies, including rabbit anti-GFRa1 (Abcam, ab186855), rabbit anti-Kit (Abcam, ab5506), rabbit anti-WT1 (Abcam, ab89901), rabbit anti-SCP3 (Abcam, ab15093), rabbit anti-γH2A.X (Cell Signaling Technology, 9718S), or rabbit anti-CLGN (Abcam, ab171971) at a 1: 200 dilution and overnight at 4 °C. After washed three times with 0.1% Triton X-100/PBS for 5 min, the cells were incubated with the secondary antibodies, including goat anti-rabbit IgG H&L (Alexa Fluor^®^ 488) (Abcam, ab150077) or donkey anti-rabbit IgG H&L (Alexa Fluor^®^ 555) (Abcam, ab150074) at a dilution with 1: 200 for 1 hour at room temperature. After washed three times with 0.1% Triton X-100/PBS for 10 min, nuclei were stained with Hoechst 33342 (Sigma, 14533-100MG) and washed three times with PBS for 5 min. The fluorescent signals were examined using a fluorescence microscope. The purity of each sample was calculated as a percentage of positive signal cells in whole cells (whole cells > 200).

### RNA extraction and RT-qPCR

Total RNA was isolated with TRIzol (Invitrogen, 15596-026) from six isolated spermatogenic cell types, including SG-A, SG-B, plpSC, pacSC, rST and eST, and then reverse-transcribed into cDNA using a PrimeScript RT reagent kit (TaKaRa, RR037A). Real-time PCR was performed on LightCycler 96 (Roche), using the SYBR Premix EX Taq kit (TaKaRa, RR820B), following the manufacturer’s protocols. mRNA expression levels were normalized to mouse Actb mRNA expression. RT-qPCR primer sequences were as follows:

Actb, Forward primer: 5′-CATTGCTGACAGGATGCAGAAGG −3′, Reverse primer: 5′-TGCTGGAAGGTGGACAGTGAGG −3′
Gfra1, Forward primer: 5′-AGGCTCAGAATTTGTTAATGG −3′, Reverse primer: 5′-TAGGGCTCAAGGGAAGGAAG −3′
Kit, Forward primer: 5′-GCCACGTCTCAGCCATCTG −3′, Reverse primer: 5′-GTCGCCAGCTTCAACTATTAACT −3′
Scp3, Forward primer: 5′-AGCCAGTAACCAGAAAATTGAGC −3′, Reverse primer: 5′-CCACTGCTGCAACACATTCATA −3′
Clgn, Forward primer: 5′-CCAGGGTGTTGGACTATGTTTG −3′, Reverse primer: 5′-CCCCGAGGAAGGTTCATCTTTA −3′

### In-situ Hi-C

The Hi-C experiments were developed following the in-situ Hi-C protocol as previously described [4] with some modifications. About 5×10^5 to 5×10^6 cells were crosslinked with final 1% formaldehyde (Sigma, F8775-500ML) for 10 min, quenched by final 125 mM Glycine (Sigma, G8898-1KG), mixed well and incubated for another 10 min. Cells were pelleted and washed once with ice-cold 1x PBS. After removing the supernatant, the cell pellets were stored at −80 °C or directly used for the following Hi-C experiments. The cells were resuspended in 500 μl of ice-cold lysis buffer (10 mM Tris-HCl, pH8.0, 10 mM NaCl and 0.2% (v/v) Igepal CA-630 (Sigma, 18896-100 ML)) containing 1x proteinase inhibitor complex (PIC, Roche, 11 873 580 001), lysed on ice for at least 20 min. After lysis, cells were pelleted at 2500g for 4 min at 4 °C then resuspended in 50 μl of 0.5% (w/v) SDS (Thermo, 24730020) and incubated at 62 °C for 10 min. Then quenched by adding 25 μl 10% (v/v) Triton X-100 (Sigma, X100-100ML) and 145 μl of water, 37 °C for 10 min with shaking (800rpm). 10 μl of MboI (NEB, R0147M) and 31 μl of 10x CutSmart buffer (NEB, B7204S) were added to the tube, then digested at 37 °C overnight with shaking. The next day, incubated the tubes at 62 °C for 20 min. Then brought volume to final 1200 μl with final 1x NEB DNA ligase reaction buffer (NEB, B0202S) and 1x BSA (NEB, B9000S), 4 μl of T4 DNA ligase (NEB, M0202L) were added and the tubes were incubated at room temperature for 4 hours with slow rotation. After Hi-C ligation, 120 μl of 10% (w/v) SDS and 20 μl of proteinase K (Thermo, EO0492) were added and incubated at 65 °C overnight. Then, the DNA were purified with ethanol precipitation and sheared to 200-500 bp by a sonicator (NingBoXinZhi, JY92-IIN). The DNA were purified again with VAHTS DNA Clean Beads (Vazyme, N411-02) and eluted in 100 μl of water. 30 μl of washed Dynabeads M-280 streptavidin beads (Thermo, 11205D) were resuspended in 100 μl of 2x Bind buffer and mixed with the purified DNA, and the tubes were incubated at room temperature for 30 min with slow rotation. After washed for 3 times, the samples were prepared for sequencing on beads. After end repairing, dATP tailing and adapter ligation, the DNA was washed 5 times with TWB buffer (5 mM Tris-HCl, pH7.5, 0.5 mM EDTA and 1 M NaCl) and resuspend in 50 μl of water. Then the DNA on beads was used for PCR amplification with phusion DNA polymerase (NEB, M0530) and purified with VAHTS DNA Clean Beads to select the DNA fragments between 200bp and 600bp. After that, all of the Hi-C libraries were sent to the company (Novogene Co., LTD) and sequenced on HiSeq X ten.

### Native ChIP-Seq

The native ChIP were performed following the previously published paper [51]. In briefly, the cells were washed once with 1x PBS and then pelleted at 800g for 5 min. The supernatant was discarded and the cells pellet was stored at −80 °C or directly used for the following ChIP experiments. For ChIP, about 2×10^5 cells were first re-suspended in 20 μl of MNase working buffer (50 mM Tris-HCl pH8.0, 1 mM CaCl2, 0.2% Triton X-100 and final 1x proteinase inhibitor complex (PIC, Roche, 11 873 580 001), and digested by 1 μl of 0.01U/μl MNase (for Histone) (Sigma, N3755-50UN) at 37°C for 2 min. 2.4 μl of 10x MNase stop buffer was then added to the tube and put on ice. Mixed with 23 μl of ice-cold 2x RIPA buffer (280 mM NaCl, 1.8 % Triton X-100, 0.2 % SDS, 0.2 % Na-Deoxycholate, 5 mM EGTA, 1x PIC) and 155 μl of ice-cold 1x RIPA buffer (10 mM Tris pH 8.0, 1 mM EDTA, 140 mM NaCl, 1 % Triton X-100, 0.1% SDS, 0.1 % Na-Deoxycholate, 1x PIC), then 16000 g at 4 °C for 10 min, and the supernatant were transferred to a new tube. Protein A/G beads (Thermo, 88803), washed with 1x RIPA buffer, were added to the reaction, 30 μl per IP reaction, incubating at 4 °C for 1 hour with slow rotation. The beads were collected via a magnet and the supernatant was transferred to a new tube. 10% were taking as Input, and 5 μl of proteinase K (Thermo, EO0492) were added and the volume were brought up to 100 μl with TE buffer (10 mM Tris pH8.0, 5 mM EDTA pH 8.0), followed by incubating with shaking at 55 °C for 1 hour and stored at −20 °C. For IP, brought volume to 100 μl with RIPA buffer for each IP reaction, mixed with CTCF (Santa Cruz Biotechnology, sc-271514/sc-28198/sc-15914) or RNA Pol II S2P (Abcam, ab5095) antibody and incubated overnight at 4 °C with slow rotation. The next day, add 30 μl protein A/G beads per IP and incubated for another 2 hours with slow rotation at 4 °C. Then washed 5 times with RIPA buffer and once with LiCl wash buffer (250 mM LiCl, 10 mM Tris-HCl pH8.0, 1 mM EDTA, 0.5% NP-40, 0.5% sodium deoxycholate), resuspended beads in 100 μl of TE buffer with 5 μl of proteinase K and incubated for 1 hour with shaking at 55 °C. For IP samples containing IgG control and the Input sample, purified the DNA and eluted in 20 μl of water. The sequencing libraries were generated by Tn5 transposase and then PCR amplification and size selected by VAHTS DNA Clean Beads. Libraries were sent for high throughput sequencing by HiSeq X Ten at Novogene.

### ATAC-seq

ATAC-seq was performed as previously described [52]. First, 5 × 10^4 cells were spun at 500 g for 5 min, 4 °C, which was followed by a wash using 50 μl of cold 1× PBS and centrifugation at 500 g for 5 min, 4 °C. Cells were resuspended in 50 μl cold lysis buffer (10 mM Tris-HCl, pH 7.4, 10 mM NaCl, 3 mM MgCl2 and 0.1% (v/v) Igepal CA-630) and centrifugation at 500 g for 10 min, 4 °C. Second, the pellet was resuspended in the transposase reaction mix (25 μl 2× TD buffer, 2.5 μl transposase (Illumina, FC-121-1030) and 22.5 μl nuclease-free water). The transposition reaction was carried out for 30 min at 37 °C. Then, the samples were purified by using a Qiagen MinElute PCR Purification Kit (Qiagen, 28004) and amplified by PCR. After size selected by VAHTS DNA Clean Beads, the samples were sequenced by HiSeq X Ten at Novogene.

### ATAC-seq data analysis

All ATAC-seq sequencing data was mapped to mm9 reference genome by snap-aligner (v 1.0) [53], the aligned reads were further sorted, indexed, removed duplicates and processed by samtools [54]. The ATAC-seq peaks of all samples were called by MACS2 using default parameter [55]. The pearson correlation coefficiency was calculated based normalized reads number on merged peaks from all samples between different samples. The ATAC-seq reads at identified peaks were quantified using bedtools [56] and normalized by using quantile norm function in R. The differential accessible chromatin regions were identified by DESeq [57]. The functional enrichment of differential accessible chromatin regions were performed using GREAT [58]. The motif enrichment analysis of differential accessible chromatin regions were performed by HOMER findMotif function [59]. The list of meiotic DNA double strand break sites was generated based on previous published DMC1 ChIP-seq [40]. The list of piRNA cluster sites was obtained from piRNA cluster database [60].

### Hi-C data mapping

The paired Hi-C sequencing reads were primary mapped, processed through HiC-Pro (v 2.8.1) [61]. First, paired reads were mapped to mm9 reference genome independently by bowtie2 and then the unmapped reads contain the MboI ligation sites were trimmed and aligned back to the mm9 reference genome again. After combing the two round mapping results, each aligned reads were assigned to one MboI restriction fragment according to the reference genome. The dangling end and self-circles pairs were excluded from valid pairs, which were further used to build the contact matrix. Most of contact matrix used in this paper were binned into 20kb size. The binned contact matrices were further normalized by using iterative correction method [61]. Further the Hi-C matrices were transferred to juicebox hic format for visualization purpose [62].

### TADs, chromatin loops and A/B Compartment Analysis

The topological associating domains (TADs) were identified as previously described, exactly following the DI HMM pipeline [3]. Briefly, the genome was divided into 40-kb windows and for each window, the frequency of interaction within 2 Mb upstream of the window to the frequency of interactions within 2 Mb downstream of the window were used to calculated the directionality index. Based on the calculated directionality index, the TADs were identified through a Hidden Markov Model. The chromatin domains were identified by using Arrowhead algorithm [4]. The chromatin loops were identified by using HICCUPS algorithm with default parameters [4]. The A/B Compartments of Hi-C 20 kb map were identified by using HOMER runHiCpca.pl. The compartment saddle analysis were performed by using cooltools (https://github.com/mirnylab/cooltools) like previously described [10,45]. To measure the strength of compartmentalization, we used the observed/expected Hi-C maps, which we calculated from 100 kb iteratively corrected interaction maps of cis interactions by dividing each diagonal of a matrix by its chromosome-wide average value. In each observed/expected map, we rearranged the rows and the columns in the order of increasing eigenvector value. Finally, we aggregated the rows and the columns of the resulting matrix into 30 equally sized aggregated bins, thus obtaining a compartmentalization plot (“saddle plot”).

### Insulation scores, aggregation TAD maps, APA and P(s) analysis

The insulation scores of each bin were calculated as previously described [63]. Briefly, to calculate the insulation score of each bin in the 20 kb binned contact matrix, the average number of interactions crossing each bin was calculated by sliding a 500 kb × 500 kb square along the diagonal of the contact matrix. The insulation score was normalized by calculating the log2 ratio of each bin’s insulation score and the mean of all insulation scores. The average insulation scores around all TADs boundaries were aggregated and plotted. To better show the dynamic change of TADs structure, the aggregated maps of chromatin interactions around TADs boundaries were plotted by basically scaling up every 2XTAD × 2XTAD chromatin maps into 100 × 100 bin matrix and plotting the average of all of these 100 × 100 matrix. The APA (Aggregation Peak Analysis) was performed as previously described [4], to measure the enrichment of loops over the local background, the KR normalized contact frequency of pixels of loops as well as the surrounding pixels up to 10 bins away in both x and y directions, i.e., 50 kb*50 kb local contact matrices, were collected and plotted. APA scores (P2LL, the ratio of the central pixel to the mean of the pixels in the lower left corner, representing the strength of chromatin loops) were determined by dividing the center pixel value by the mean value of the 25 (5*5) pixels in the lower right section of the APA plot. The P(s) analysis calculated the relationship curve between chromatin interaction probability and linear distance between two DNA fragments as previously described [10,49]. Briefly, contact probability (P(s)) curves were computed from 100 kb binned Hi-C data, and we divided the linear genomic separations into logarithmic bins with a factor of 1.3. Data within these log-spaced bins (at distance, s) were averaged to produce the value of probability.

### ChIP-Seq analysis

The Pol II ChIP-Seq data [64], our Pol II S2P ChIP-Seq data and CTCF ChIP-Seq data were mapped to mm9 reference genome using snap-aligner (v1.0) [53]. And the ChIP-Seq peaks of CTCF were called by MACS2 with default parameters [55]. The CTCF binding ratio near TADs boundary was calculated by dividing CTCF peak numbers in each distance bin by total CTCF peak numbers. All of the visualization of reads were processed on IGV genome browser [65].

## Supporting information

**S1 Fig. Purities of eight isolated spermatogenic cell types. (A)** The schematic of mouse spermatogenesis (Top panel). Phase-contrast microscope showed the morphological characterization of eight isolated spermatogenic cell types, including PriSG-A, SG-A, SG-B, plpSC, pacSC, rST, eST and SZ (Bottom two panels). Scale bar, 50 μm. **(B)** Cell purities according to the immunofluorescence data and our published results[27], the total cell number for each sample was more than 200. **(C)** Immunofluorescence revealed the expression of germ cell type-specific marker proteins in four isolated spermatogenic cell types and sertoli cell (SE). Specifically, GFRa1 for PriSG-A, Kit for SG-A, SCP3 and γH2A.X for pacSC, CLGN for rST and WT1 for SE. Scale bar, 10 μm. **(D)** Expression levels of the male germ cell specific marker genes mRNA transcripts (Gfra-1 for PriSG-A, Kit for SG-A, Scp3 for pacSC, and Clgn for rST).

**S2 Fig. High correlation of Hi-C data and chromatin accessibility during spermatogenesis by ATAC-seq. (A)** P(s) analysis of six stages’ independent replicates of Hi-C data suggested high correlation between biological replicates. **(B)** The scatter plot of two replicates’ of priSG-A, SG-A, pacSC and rST ATAC-seq signals. **(C)** Metagene plot of ATAC-seq signals at promoter regions of known genes showed that chromatin accessibility was gradually decreased during spermatogenesis. **(D)** Metagene plot of ATAC-seq signals at enhancer regions showed that chromatin accessibility was gradually decreased during spermatogenesis.

**S3 Fig. The landscape of chromatin accessibility during spermatogenesis. (A)** The quantification heatmaps of all chromatin accessible sites identified in four cell types of spermatogenesis. **(B)** The functional enrichment of the differentially accessible chromatin regions between PriSG-A and pacSC. **(C)** The functional enrichment of the differentially accessible chromatin regions between pacSC and rST. **(D)** The identified transcription factor (TF) motifs enriched in closed chromatin regions between PriSG-A and pacSC. (i. e. closed in PriSG-A and opened in pacSC). **(E)** The identified transcription factor (TF) motifs enriched in opened chromatin regions between PriSG-A and pacSC. (i. e. opened in PriSG-A and closed in pacSC). **(F)** The identified transcription factor (TF) motifs enriched in closed chromatin regions between pacSC and rST (i. e. closed in pacSC, and opened in rST). **(G)** The identified transcription factor (TF) motifs enriched in opened chromatin regions between pacSC and rST. (i. e. opened in pacSC, and closed in rST). **(H)** The number of accessible DSB sites between PriSG-A and pacSC. **(I)** The number of accessible piRNA clusters between PriSG-A and pacSC.

**S4 Fig. A/B compartments still existed at pacSC and were dynamic during spermatogenesis. (A)** The percentage of genome-wide active and inactive genomic regions at PriSG-A, SG-A, SG-B, plpSC, pacSC, rST, eST and SZ stages during spermatogenesis based on compartments. **(B)** Compartmentalization saddle plots: average distance-normalized interaction frequencies between cis-pairs of 100-kb bins arranged by their eigenvector value (EV1).

**S5 Fig. The A/B compartments switch during mouse spermatogenesis. (A)** The dynamic changes of PC1 values (Compartments) during spermatogenesis. **(B)** The correlation between PriSG-A and other stages during mouse spermatogenesis based PC1 values (Compartments). **(C)** The percentage of A to B transitions and B to A transitions from PriSG-A to SG-A, SG-A to SG-B, SG-B to plpSC, plpSC to pacSC, pacSC to rST and rST to SZ. **(D)** The compartment changes at the genomic locus of Kit gene.

**S1 Table. The quality control of Hi-C data.**

## Acknowledgements

We sincerely thank Dr. Mingxi Liu, Dr. Bo Wen, and Dr. Yong Zhang for their helpful advice.

## Author contributions

### Conceptualization

Fei Sun, Xiaoyuan Song

### Formal Analysis

Ruoyu Wang, Yusheng Chen

### Funding Acquisition

Fei Sun, Xiaoyuan Song

### Investigation

Zhengyu Luo, Xiaorong Wang, Jian Chen, Qianlan Xu, Jun Cal, Xiaowen Gong

### Methodology

Zhengyu Luo, Xiaorong Wang, Ruoyu Wang

### Project Administration

Fei Sun, Xiaoyuan Song

### Resources

Ji Wu, Yungui Yang, Chunsheng Han, Fei Sun, Xiaoyuan Song

### Software

Ruoyu Wang, Yusheng Chen

### Supervision

Fei Sun, Xiaoyuan Song

### Writing – Original Draft

Zhengyu Luo, Xiaorong Wang, Ruoyu Wang, Xiaoyuan Song

### Writing – Review & Editing

Zhengyu Luo, Xiaorong Wang, Ruoyu Wang, Wenbo Li, Fei Sun, Xiaoyuan Song

## Data availability

The raw sequence data reported in this paper have been deposited in the Genome Sequence Archive [66] in the BIG Data Center [67], Beijing Institute of Genomics (BIG), Chinese Academy of Sciences, under accession numbers CRA001095 that is publicly accessible at http://bigd.big.ac.cn/gsa.

